# Spatial trajectories of coffee harvesting in large-scale plantations: ecological and management drivers and implications

**DOI:** 10.1101/2023.10.31.564942

**Authors:** Emilio Mora Van Cauwelaert, Denis Boyer, Estelí Jiménez-Soto, Cecilia González González, Mariana Benítez

## Abstract

**CONTEXT:** Coffee is produced under different management systems and scales of production categorized as Syndromes of Production. The “Capitalist Syndrome” is characterized by the high use of capital and labor inputs to increase agricultural outputs. This syndrome results in practices like high planting densities that may promote the development and dispersal of plant pathogens like coffee leaf rust. The spatial arrangement of coffee trees drives the spatial movement of the harvesters, who can bear and disperse pathogens across and within plantations. In most capitalist coffee plantations, harvesters work multiple hours to maximize the daily harvest, which might increase their dispersal potential. However, their spatial movement has not yet been described, nor its relationship with the scale or management of the plantation, and even less its ecological implications for pathogens dispersal.

**OBJECTIVE:** We describe and analyze the daily spatial movement of coffee harvesters in two large-scale capitalist plantations: an organic and a conventional plantation.

**METHODS:** Using state-space models, we recorded and analyzed the spatial movements of harvesters. We then constructed a driver tree for harvest dynamics, which incorporated qualitative variables related to climate, coffee plants, and management aspects reported by the harvesters.

**RESULTS AND CONCLUSIONS:** Our model differentiated two kinds of movements: 1) when trees have berries, harvesters remain in the coffee rows or areas nearby (Collect state; 94-98% of the steps); 2) when not, harvesters make longer steps within the harvesting location or move to another area (Search state; 2-6% of the steps). In the organic plantation, the Search state had a longer-tailed step-length distribution than in the conventional plantation, resulting in a significantly higher visited area per worker (p<0.05). This might be related to a) a lower fruit load or percentage of trees with ripe fruits when we took the data or b) smaller harvesting locations (“*pantes*”) per number of harvesters. Harvesting movements that explore a wider area, either by visiting more plants or by changing locations on the same day, could create more foci of CLR infection across the plantation.

**SIGNIFICANCE:** Our results highlight practices that can reduce the possible impact of human dispersal of pathogens like shorter harvesting trajectories by working fewer hours a day or avoiding harvesting at the end of the maturation season when few trees have berries and harvesters have to travel longer distances. This calls for organic coffee management that could prevent diseases, increase diversity, and guarantee just and safe conditions for workers.

## 1. INTRODUCTION

In recent decades, the increased industrialization of agroecosystems has gathered significant attention due to its profound negative social and ecological consequences (Raven and Wagner, 2021). Agricultural industrialization and environmental change have led to increases in pest populations (Andow, 1983), biodiversity decline (Raven and Wagner, 2021), and are also associated with a higher likelihood of zoonotic disease outbreaks (Jones et al., 2013; Perfecto et al., 2023). Adding to these challenges are the heightened demands on human labor and precarious social conditions experienced in large-scale industrial agricultural systems (Mapes, 2010), a social phenomenon that imparts more complexity to agroecosystem management and processes.

Coffee agroecosystems exhibit a large variety of management practices that reinforce each other and that can be defined as “syndromes of production”, a conceptual framework useful to evaluate social and ecological dynamics across distinct agricultural paradigms (Andow and Hidaka, 1989; Moguel y Toledo, 1999; Ong and Liao, 2020; Vandermeer and Perfecto, 2012). Coffee production syndromes can be mainly distinguished by shade management (shade or sun coffee plantations; Vandermeer and Perfecto, 2012) or by the type of land tenure and scale of production (large-scale capitalist or small-scale peasant-owned coffee plantations; Ong and Liao, 2020). In this sense, coffee agroecosystems are model systems to understand how different syndromes of production affect ecological processes such as their planned and associated biodiversities (Mas and Dietsch, 2004) but also social aspects like the worker’s conditions (Jiménez-Soto, 2021).

The capitalist syndrome follows a maximization of profit rule (Ong and Liao, 2020); it is typically based upon large-scale and high-density planting farms with a high input of human labor and exploitation, although it may have specific ecological practices according to organic or conventional coffee markets (Perfecto et al., 2019). This syndrome of coffee production results in practices that may promote the development and spread of important pests (Avelino et al., 2006; Hajian-Forooshani et al., 2016). For instance, the coffee leaf rust (CLR) can travel across and within the coffee plantations through direct leaf contact, water splash, or local turbulent wind conditions (Becker and Kranz, 1977; Vandermeer et al., 2018), but its effectiveness of dispersal and impact can be modulated by management practices like the planting density (Mora Van Cauwelaert et al., 2023), the presence of wind-barrier trees in the plantation (Gagliardi et al., 2020), or the biotic communities of natural enemies (Jackson et al., 2012), all determined to some degree by the syndrome of production (Vandermeer and Perfecto, 2012; McCook and Vandermeer). In this regard, increased agricultural simplification and industrialization of coffee agroecosystems have led to massive outbreaks of CLR around the globe (Avelino et al., 2015; McCook and Vandermeer, 2015).

Some studies have proposed that harvesters can also bear CLR spores during harvesting (Becker and Kranz, 1977; Waller, 1981). Interestingly, the range of movement and duration of the daily harvest are also determined by the management, the scale, the land ownership, and by the agricultural characteristics of the coffee plantations. In most landlord-owned large-scale coffee plantations, harvesters are paid by piecework and work several hours to maximize the daily harvest (Jiménez-Soto, 2021; López Echeverría, 2006). These practices affect the social and health conditions of the workers, which are often precarious. Besides, characteristics like high planting density can also drive the spatial movement during harvesting. We argue that the spatial movement of the harvesters under capitalist labor-intensive conditions could also have an ecological impact, through modulating the human dispersal of pathogens like coffee leaf rust (CLR). In fact, other studies have stressed the relation between the landscape structure, individual movement behavior, and pathogen transmission for predicting and understanding disease dynamics (Manlove et al., 2022; White et al., 2018).

However, the spatial movement of the harvesters in plantations within this capitalist syndrome of production has not been described yet, nor its relationship with the management of the plantation, and even less its ecological implications for CLR or other potential pest dispersal. Here we present a description and qualitative analysis of the spatial movement of farmworkers during harvesting in two large-scale landlord-owned plantations (∼300 ha) with two different types of management: an organic shaded production coffee plantation and a sun-grown production with few scattered trees. Specifically, we asked 1) how to statistically characterize the spatial trajectories of workers during harvesting, 2) how different features of the plantations, determined within a specific syndrome of production, affect these trajectories, and 3) what are the socio-ecological consequences of the harvesting movement in these large scale plantations.

We first recorded and analyzed with state-space models the daily spatial movements of the harvesters, as well as their differences between both plantations. We then constructed a driver tree for harvest dynamics, which incorporated qualitative variables related to climate, as well as coffee biology and management aspects reported by the harvesters during interviews and noted through our own observations. We also highlighted the relationship between these variables and the syndrome of production. Finally, we put in conversation the patterns and dynamics of the spatial movement of harvesters with the social conditions associated with the large-scale capitalist syndrome and the possible implications in pathogen dynamics such as CLR spread.

## 2. METHODS

### 2.1. Site of study

We selected two large-scale coffee plantations located in the Soconusco region of Chiapas, Mexico. This region has a tropical wet and dry climate (Koppen classification, Aw), with a rainy season from May to October, an annual mean temperature of 27 °C, and a precipitation of 2800 mm (Villarreal-Treviño et al., 2015). The first one (organic plantation, henceforth) is an organic shaded coffee plant plantation with a high plant diversity of up to 200 species. The second one (the conventional plantation) has a lower shade cover dominated by few *Inga sp*, uses pesticides, and has a high rate of coffee tree renovation to increase the productivity of regions affected by coffee leaf rust (CLR). Both plantations are approximately 300 ha, are landlord-owned and produce different coffee varieties for exportation. Harvesting is carried out by permanent or temporary workers from Central America, primarily Guatemala and El Salvador, from October to February, depending on the previous rainy season. Workers are paid by piecework, either by weight or volume, and live in precarious conditions (Jiménez-Soto, 2021). During harvest, the foreman assigns the harvesters and their families specific areas called “*pante*” (a Nahuatl word referring to agricultural locations of variable size) to work during one or multiple days. Harvesters remain in these locations during the day or travel to another area when they finish harvesting all the trees in the *pante*.

### 2.2. Data Collection

We carried out field research for three weeks between October and November of 2021. We accompanied 12 families of harvesters during their harvesting day, six on each plantation, after their explicit consent to our research. The trajectories were labeled from C1 to C6 and from O1 to O6 for the conventional and organic plantations, respectively. We worked with them for six hours (including resting and eating pauses), completing three tasks: i) We recorded the movement of one harvester from each family by marking the location of every newly harvested tree, ii) we carried out semi-structured walking interviews (Evans and Jones, 2011) with each family and participant observation during harvesting to assess the factors that determined the different trajectories and to gather information about the whole syndrome of production, and iii) we participated in harvest, as one team member acquired the necessary harvesting skills along with experienced harvesters and donated their daily harvest to contribute to the family process.

#### 2.2.1. Trajectories data

We added a waypoint for each consecutive visited tree using the satellite remote sensing Global Positioning System (GPS) Garmin 65. This system is ideal to analyze fine-scale movement patterns as its relative distance error between two consecutive recorded points is small (<0.5 m) (Breed and Severns, 2015). The recorded tracks were homogenized and curated by removing the pauses and normalizing the UTM coordinates. We defined the step length as the distance between two harvested trees; this distance provides a good proxy of the general movement of harvesters and is also relevant to understand how patterns of movement could explain the dispersal of CLR or other potential pests from one tree to another. Every pair of trees closer than one meter was considered as the same point (Vandermeer et al., 2018).

#### 2.2.2 Walking interviews

We designed a questionnaire to gather information on the organization of harvest and applied it as a walking interview with the harvesters (Evans and Jones, 2011). We included questions about the period of the harvesting season, the number of times each plot was harvested, and the differences in the harvesting dynamics along the season between both plantations (organic and conventional). In particular, we asked who decided the size of the groups, the starting point, and the daily length of harvest. We also asked the harvesters directly about the factors that affected the motion of harvesting within each plant and between the plants. We added questions about the CLR control in both plantations and, last but not least, about the origins and social conditions of the harvesters. Before this process, we explained the purpose of the study and gave an informative letter to each of the participants to obtain their informed consent to participate in the study. The data was complemented with personal observations on the characteristics and times of harvesting per plant, the size of the harvesting groups, and the differences in topography and ecological management between both plantations like the planting pattern of coffee trees.

### 2.3. Movement analysis and qualitative drivers of motion

We analyzed the harvester movement data using state-space models (Patterson et al., 2008) and organized the factors that influence the motion in a qualitative driver tree. All data analyses and figures were done in *Rstudio* 2023.03.1+446 using *plyr, dplyr, tidyverse, ggplot2, patchwork* libraries, and *Inkscape 1.0 and yEd 3.23.2*. All code and data required to reproduce results in the work can be accessed at https://github.com/tenayuco/harvestDistribution and it is further explained in supplementary material.

#### 2.3.1. Movement analysis, state-space models and differences between the plantations

We analyzed the harvesters’ movement using multi-state distribution models. State-space models are typically used in the context of movement ecology analysis to characterize the trajectories of individual animals that are assumed to have different behaviors or states (eg. foraging, exploring, migrating) during the recorded movement (Michelot and Blackwell, 2019; Morales et al., 2004; Patterson et al., 2008). These behaviors are derived by characteristics like the relative angle between two steps, or the step length. We used this approach to statistically quantify differences in the daily movement of the harvesters (for the complete details, refer to Supplementary Material).

We first fitted the step length frequency distributions of the six recorded trajectories of each plantation to models of one and two-states distributions (i.e. models assuming that the observed movements are generated by one or by two behavioral states during harvest). In the two-state case, we termed the behaviors as “Collect” and “Search”. In all cases, we used two different fitting distributions, namely, the Gamma (G(x)) and Weibull (W(x)) functions (see Equations (1-3); Fig.S1.3; Table S1.2). For ***x*** and ***α >0***, these normalized probability density functions (PDFs) are given by the following formula:

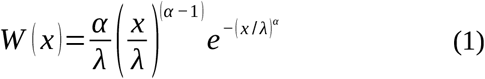

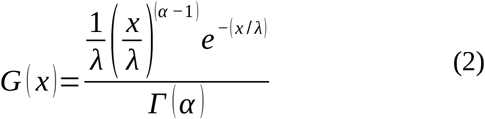

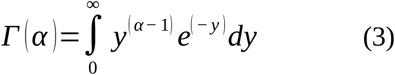

where ***x*** is the step length (in m), ***α*** is a shape parameter (no units), ***λ*** a scale parameter (in m) and ***Γ(α)*** the gamma function (Pishro-Nik, 2014). In the two-state case, the Search state should typically be characterized by a significantly larger ***λ*** than the Collect state. In each fitting process, we used the forward algorithm implemented in the *movehmm* library (Michelot et al., 2016) with 1000 combinations of prior parameters from uniform distributions that included the mean and the maximum step lengths of the data (Table 1). For the analysis, the steps were time-irregular and we assumed no differences between the harvesters. We compared the Gamma and Weibull models, as well as the one-state and two-state models, using the Akaike (AIC) criteria (Michelot et al., 2016). As the two-states distribution of the Gamma family (equation 2-3) generated a lower AIC for both plantations, we chose it to assign to each step of the trajectories one of the two states (Search and Collect), using Hidden Markov Models (HMM) and the Viterbi algorithm (Zucchini et al., 2016). Hidden Markov models (HMMs) assume that the distribution that generates an observation Zt (a step-length) depends on the state St of an underlying and unobserved Markov process. In brief, this procedure jointly fits the two-state distribution of step-lengths and establishes a sequence of states using a maximum likelihood procedure (Patterson et al. 2008; Zucchini et al., 2016). It first assigns to each step the most likely state and then corrects the state using a global decoding procedure with the whole sequence (Zucchini et al., 2016). Finally, we explored the differences in the obtained distributions for each state (Search and Collect) between both plantations and their implications in the number of trees and surface area visited. We divided the total area into squares with different sizes (from 1 to 40 m wide) to account for the effect of the grain size and assumed that a square was visited if at least one visited tree fell within it. We tested the significance of the difference between the means for each measurement using the Wilcoxon test (w) or the Student test (st) depending on the normality of the residuals.

**Table 1.**
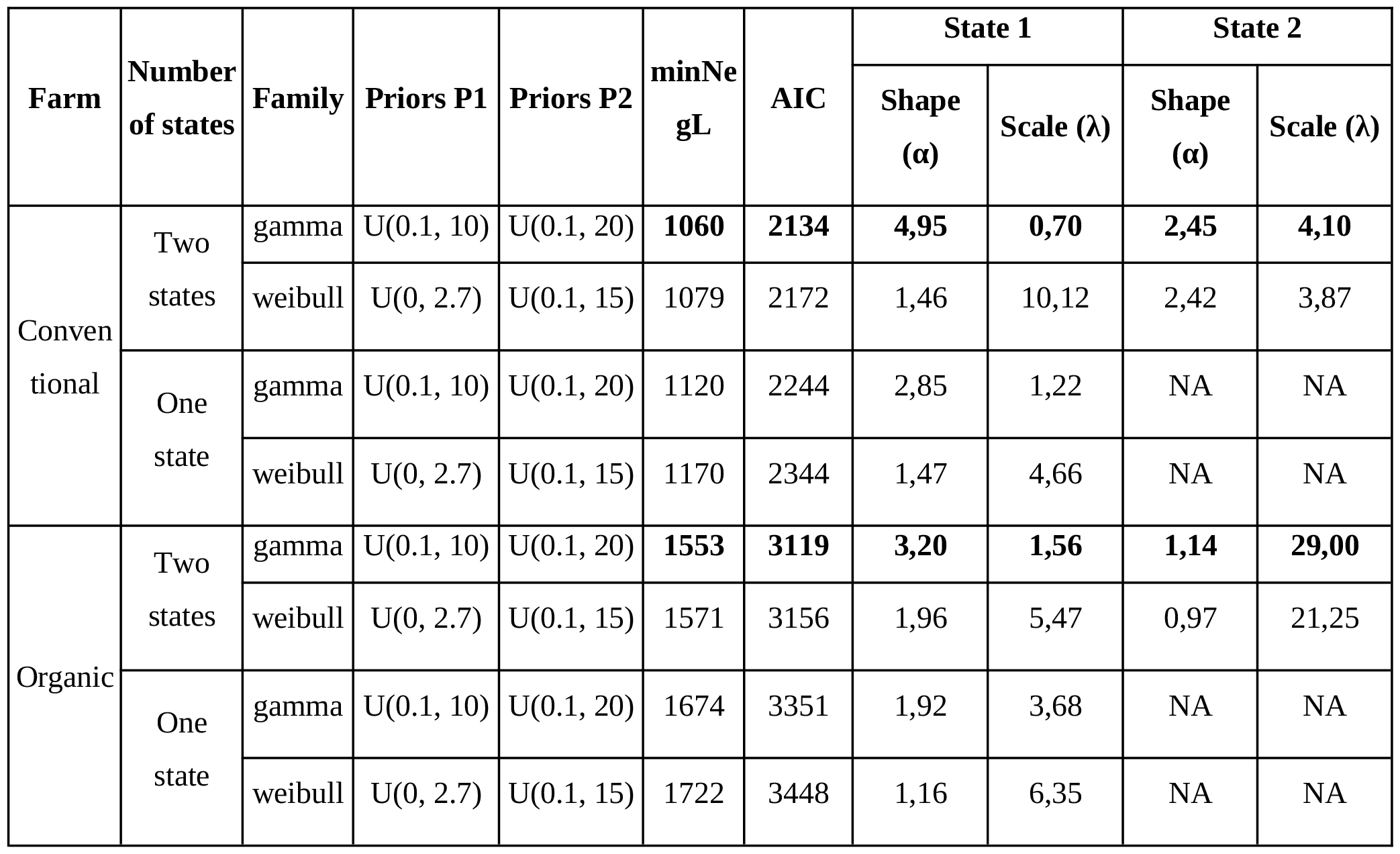
Best models for each plantation and distribution family, for one and two-state models (State 1=Collect, State 2=Search). We added the prior parameters for parameter P1 (mean and shape for gamma and weibull, respectively) and P2 (standard deviation and scale for gamma and weibull, respectively) (Michelot et al., 2016). We show the minimal negative likelihood (minNegL), AIC values and the final parameters obtained and transformed to shape and scale following the following relations: α = µ^2^/σ^2^, λ= σ^2^/µ, where α:shape, λ:scale, µ:mean, σ:standard deviation. The parameters in bold are the ones we used for the analysis (equations (2)-(3)).

| Farm | Number of states | Family | Priors P1 | Priors P2 | minNegL | AIC | State 1 |  | State 2 |  |
| --- | --- | --- | --- | --- | --- | --- | --- | --- | --- | --- |
| | | | | | | | Shape ($\alpha$) | Scale ($\lambda$) | Shape ($\alpha$) | Scale ($\lambda$) |
| Conventional | Two states | gamma | U(0.1, 10) | U(0.1, 20) | **1060** | **2134** | **4,95** | **0,70** | **2,45** | **4,10** |
|  |  | weibull | U(0, 2.7) | U(0.1, 15) | 1079 | 2172 | 1,46 | 10,12 | 2,42 | 3,87 |
|  | One state | gamma | U(0.1, 10) | U(0.1, 20) | 1120 | 2244 | 2,85 | 1,22 | NA | NA |
|  |  | weibull | U(0, 2.7) | U(0.1, 15) | 1170 | 2344 | 1,47 | 4,66 | NA | NA |
| Organic | Two states | gamma | U(0.1, 10) | U(0.1, 20) | **1553** | **3119** | **3,20** | **1,56** | **1,14** | **29,00** |
|  |  | weibull | U(0, 2.7) | U(0.1, 15) | 1571 | 3156 | 1,96 | 5,47 | 0,97 | 21,25 |
|  | One state | gamma | U(0.1, 10) | U(0.1, 20) | 1674 | 3351 | 1,92 | 3,68 | NA | NA |
|  |  | weibull | U(0, 2.7) | U(0.1, 15) | 1722 | 3448 | 1,16 | 6,35 | NA | NA |

#### 2.3.2. Qualitative drivers of the spatial trajectories

To understand the qualitative drivers of harvesters’ trajectories we organized the variables obtained during the walking interviews to analyze their relation with the longitude, the direction of the trajectories, or the presence of short and long tailed step-length distributions. We separated the variables into two categories: environmental or coffee biology-related (e.g. rainfall or coffee interplant ripening percentage) and those that could be directly affected by management decisions or, in general, by the syndrome of production (e.g. coffee variety, size of the *pantes*). Finally, we connected these variables when they influenced each other and categorized the connection into direct and indirect drivers. Direct drivers included those whose modification affected the trajectory. Indirect factors affected direct factors (but not the trajectory itself).

## 3. RESULTS

### 3.1 The movement of the harvesters can be described by two-state distribution models with qualitative differences between the plantations

Harvesters visited 94.5 trees on average (± 25.6) during the three to four hours recorded daily, excluding pauses. The average time between each visited tree was 2.3 min (± 1.8 mins). The spatial trajectories of the 12 harvesters were largely determined by the planting pattern of coffee trees (Fig.1). Within these patterns some harvesters followed more or less specific coffee tree rows (C_1 to C_3, C_5, O_6) or jumped from one row to another or other regions, depicting more randomized trajectories (C_4, C_6, O_2 to O_5) (Fig. 1). The aggregated distribution of the step lengths of harvesters in each plantation (*n*=6) is represented in Fig. 2. In both plantations the distribution is skewed to the right. In the conventional plantation the mean step length is 4.2 m and the median is 3.2 m whereas in the organic plantation these values are 6 m and 4.5 m, respectively. In some cases, harvesters could travel up to 50 m from one tree to another (conventional plantation), or even more than 100 m (organic plantation) (see inset boxplots in Fig.2).

**Figure 1.**
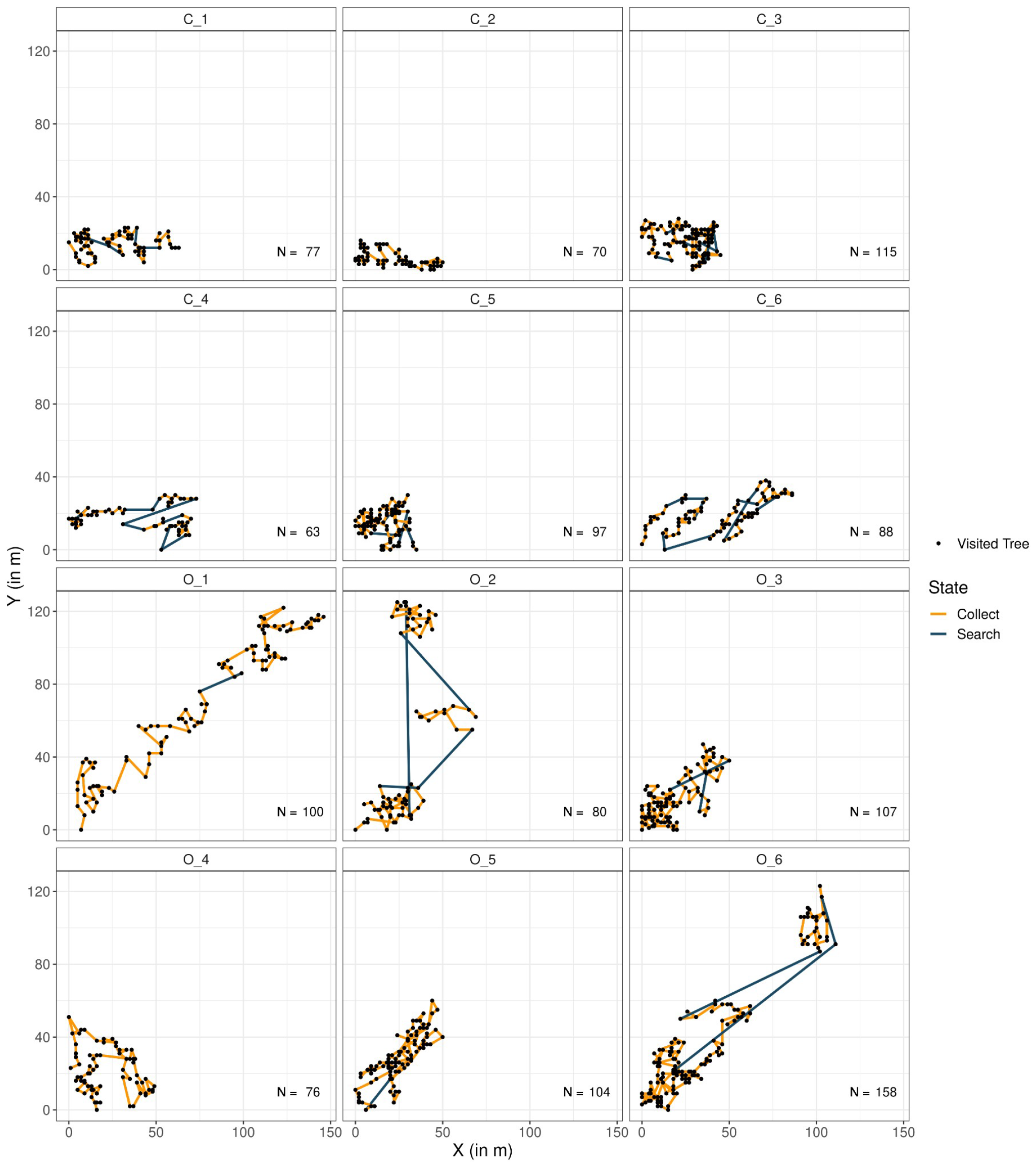
Harvesting trajectories from the conventional (C) and from the organic (O) plantations. We normalized the UTM coordinates from 0 to 150 m to represent the trajectories. The colors represent the most likely state to which each step belongs for each of the farms (orange: Collect, dark blue: Search; see the text for a full explanation). Each black point represents one harvested tree.

**Figure 2.**
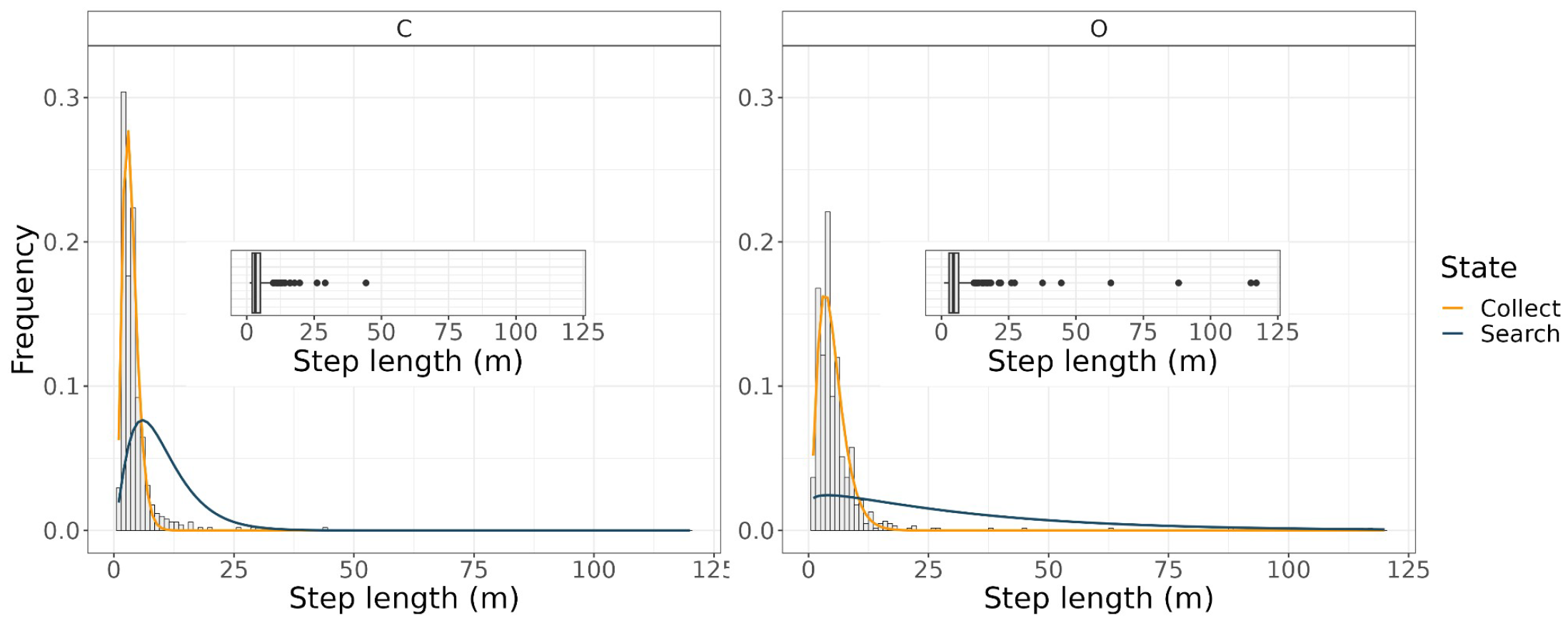
Histograms and box plots (insets) of the step lengths for the trajectories in each plantation and the probability density function of the gamma distributions for each of the two states. In the conventional plantation (C), the parameters of the gamma distribution (α, λ) are (4.95, 0.7) in the Collect state (orange line) and (2.45, 4.10) in the Search state (dark blue line). In the organic plantation (O) these parameters become (3.2, 1.56) for the Collect state t; and (1.14, 29) for the Search state . See Table 1 and equations (2)-(3).

Models with two distinct state distributions provided a better fit to the aggregated step length distribution as they exhibited lower general AICs, for both Gamma and Weibull distributions, than models with only one state (Table 1 and Fig. S1.5). In particular, the Gamma family with two states had the lowest AIC in both plantations (Table 1). Each step along the trajectories was assigned a state in Fig. 1 according to the distributions of Fig.2. In our work, the states are interpreted as a proxy for the situation during harvest. We named the first state “Collect”, where the worker remains in the same vicinity of the *pante* harvesting nearby plants (Fig.1). The PDF of this state is qualitatively similar in both plantations (Table 1, Fig. 2) and encompasses steps from 1 to 8.6 m in the conventional plantation (94% of the steps) and from 1 to 18 m in the organic plantation (98% of the steps) (Fig.2). In the second state, the “Search” state, harvesters traveled to another zone of the *pante* or to another *pante* when the current one had been completely harvested (Fig. 1). This second Search state is described by two qualitatively different PDFs depending on the plantation (Table 1, Fig.2). In the conventional plantation, it includes steps from 7.4 to 44 m and represents movements within the *pante* (6% of steps, dark blue line in Fig. 1 and Fig.2). In the organic plantation, this state includes steps from 21 to 117 m long and represents mainly movements between *pantes* or relocalizations (2% of steps, dark blue line in Fig. 1 and Fig.2). Also note that in the organic plantation, the shape parameter α is close to 1 in the Search state (exponential distribution) while it is above 3 in the Collect state (distribution with a maximum at a finite step length). In the conventional plantation, the difference between the two states is less pronounced.

### 3.2. Harvested trees and visited area per day are higher in the organic plantation

The average distance between trees within each *pante* is only slightly larger in the organic plantation (C: 3.8 ± 2.11 m; O: 4.8 ± 2.6 m; not significant). In this sense, the marked differences in the PDFs distributions of Search and Collect states between both plantations are not completely determined by the planting pattern. The average number of trees visited per unit of time were higher in the organic than in the conventional plantation but the differences were not significant (C: 85 (± 19) trees per day, O: 104 (± 29) trees per day; C: 25 (± 7) trees per hour; O: 29 (± 5) trees per hour) . The higher number of harvested trees did not seem to correspond to differences in harvest weight per worker, which implies that, in the organic plantation, each tree had less mature berries when the data was taken.

The average traveled area per worker (approximated as the number of visited squares of given side length) was significantly higher in the organic plantation, for square units of 3, 5, 10 and 20 m side (Fig. 3). When the unit area is too small (1 m^2^) or when the unit is too large (1600 m^2^ or 40 m side) no significant differences are observed. The higher visited surface area might be the consequence of a higher presence of long steps (or relocalizations) during the harvesting day (Fig. 1 and Fig. 2). In some specific trajectories harvesters traveled up to 100 meters to reach the following tree.

**Figure 3.**
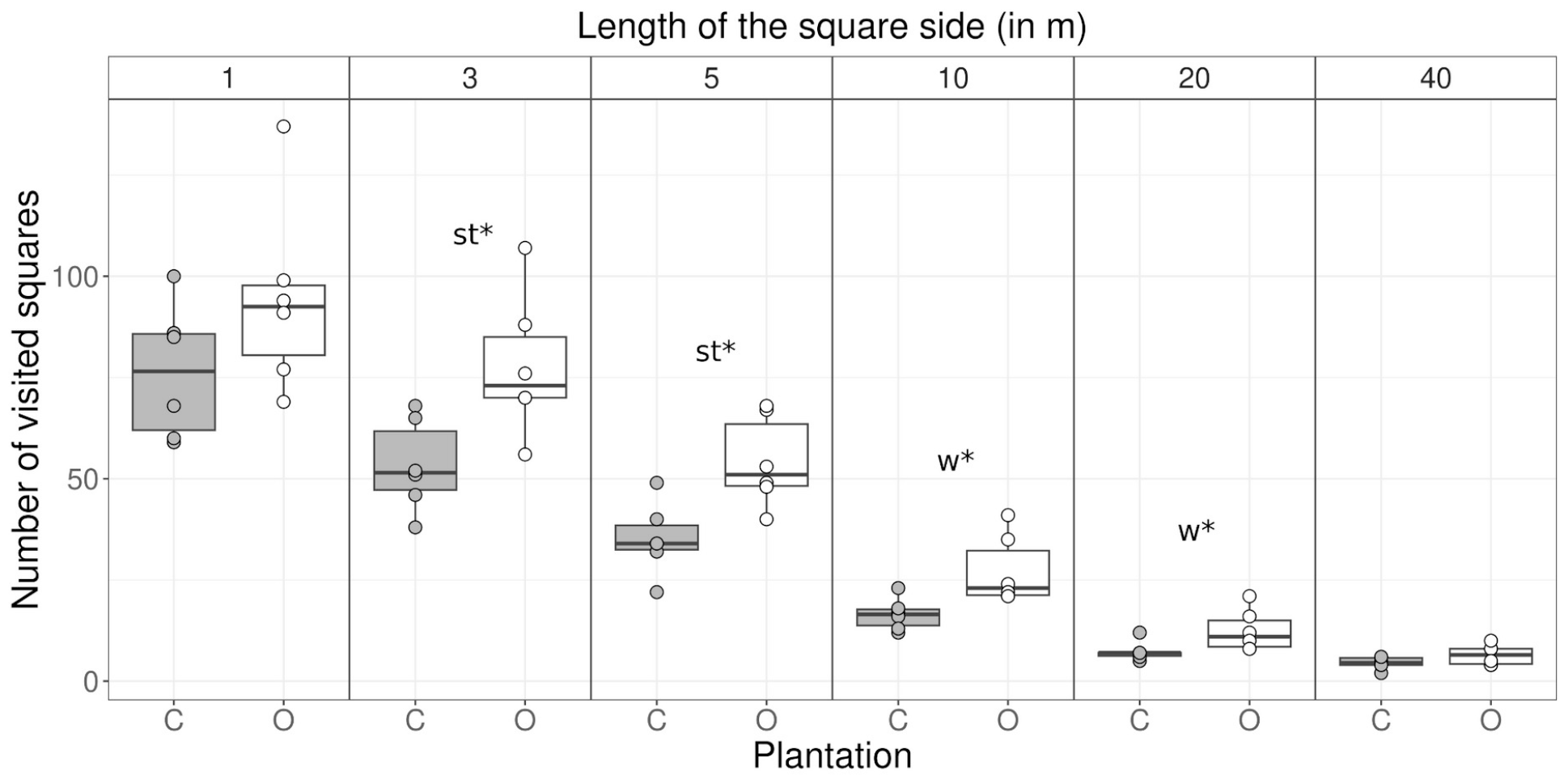
Surface area visited per harvester, obtained from visited squares of different sizes. Each dot represents the number of visited squares per harvester in each of the plantations (C-gray: conventional, O-white: organic). For the square unit areas where the differences between the two plantations were significant we added a ^*^ (p<0.05) and the used test of comparison between the two means depending on the normality of the residuals (st: Student, w: Wilcoxon, n= 6).

### 3.3. Differences in trajectories of the harvesters are mediated by climate, the coffee biology and plantation variables, and most of them are modulated by management decisions

The spatial movement of the harvesters, in particular the presence of longer steps (mainly in the state Search), as well as the daily length of the trajectory, can be directly and indirectly mediated, at lea st, by climate, topography, coffee biology characteristics, and coffee management variables (Fig. 4; Table 2). The relationship between these variables give us some insights on the determinants of the syndrome of production and the reasons that might explain the differences and commonalities in the trajectory between both plantations.

**Table 2.**
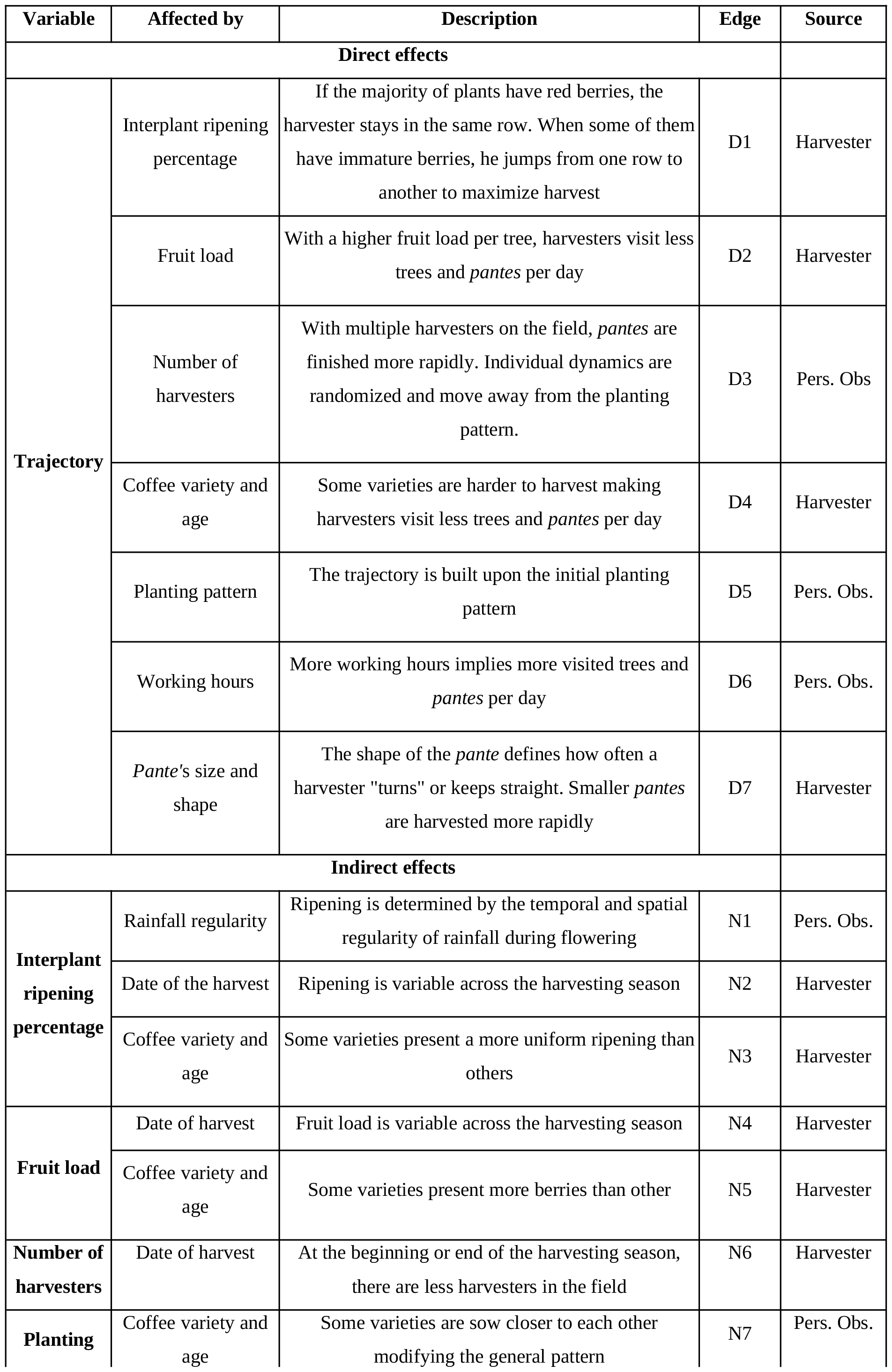
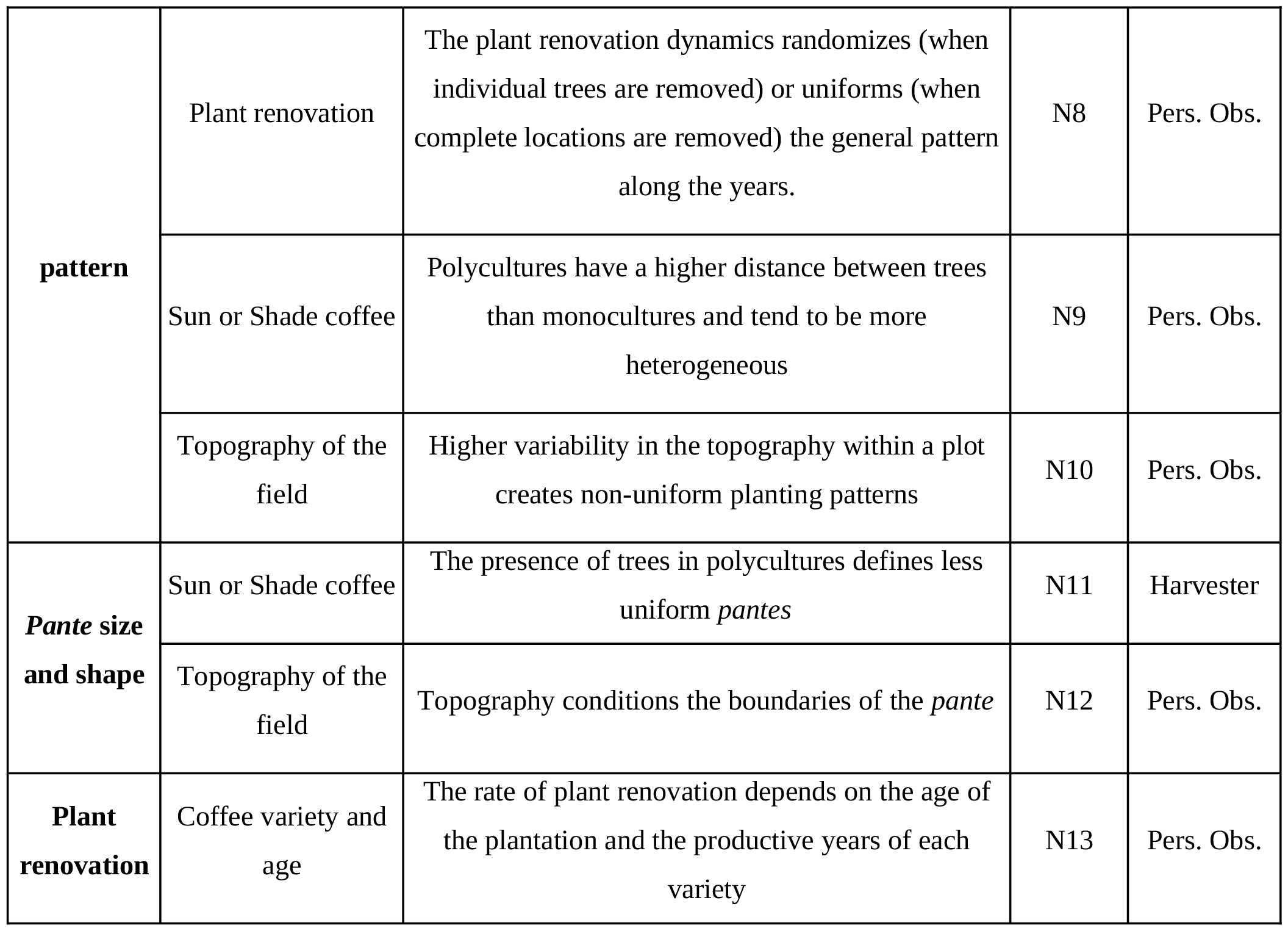
Direct (D) and indirect (N) drivers of the trajectories of the harvesters. The relation between the nodes is shown in Fig. 4.

**Figure 4.**
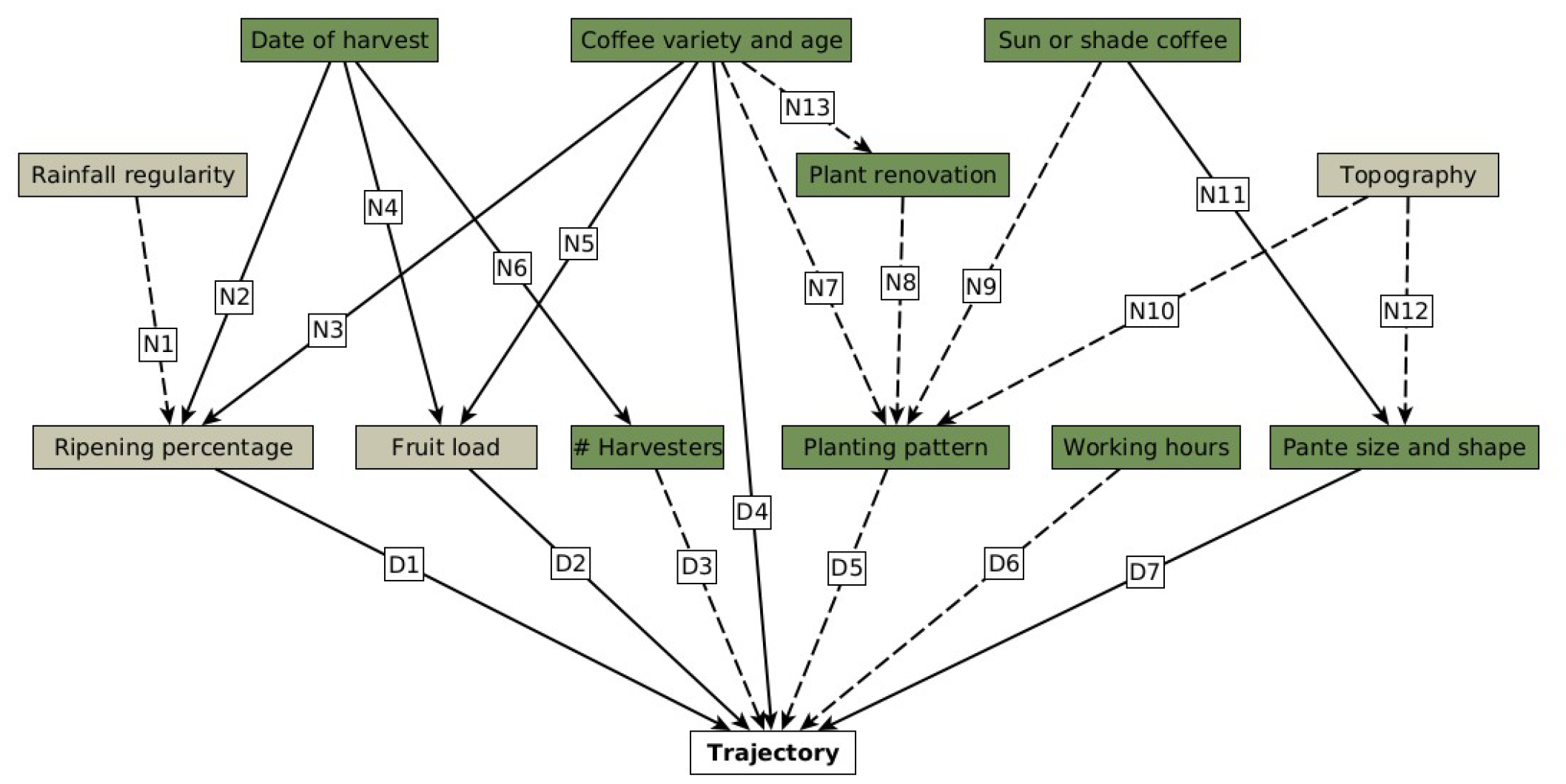
Qualitative drivers of the spatial trajectories of the harvesters during harvesting. The nodes represent the variables that interplay with the trajectory and the edges the causal relationships between them. The color of the boxes indicates if the variable is environmental or coffee biology related (gray), and if it can be directly modified by the management of the plantation (green). Note that the modification of these variables affects all the variables downstream. The letter in the label of the edges indicates whether the effect on the trajectory is direct (D) or indirect (N). The explanation of each label is summarized in Table 2.

According to the interviews and our additional observations (solid and dotted lines in Fig. 4, respectively), the harvester’s trajectory might be directly affected by the number of mature berries in each of the trees. This number is determined by the fruit load (total number of berries per tree) and the proportion of mature berries per tree. Here we use “fruit load” to refer to the number of mature berries per tree (D2). When trees have more berries, harvesters visit fewer trees and *pantes* per day, and their final trajectory is shorter (Table 2). Besides, to maximize their daily harvest, harvesters keep the same row when all the trees have berries, or jump from one row to another or even to another *pante*, when some trees are still immature. We refer to this as the interplant ripening percentage (i.e. the percentage of trees with ripe berries, D1). Low ripening percentage results in more randomized patterns or larger step lengths along the day (state Search) (Fig.1). Ripening percentage and fruit load depend on the coffee variety and age (N3, N5) and the date of harvest (N2, N4). Around the middle of the season, each coffee variety has a peak of berry maturation and synchronization between the trees. When we collected our data, trees were more homogeneously loaded in the conventional plantation than in the organic plantation, explaining in part the higher number of harvested trees in the organic plantation and the presence of larger step-lengths in the state Search (Fig. 1). This pattern could have been reversed in another period. It is worth mentioning that coffee ripening is mostly mediated by the rainfall regularity during the flowering season (N1) and the coffee variety (N5).

On the other hand, the spatial movement of harvesters is built upon the initial planting pattern (in rows, in groups, etc) and the *pante* size and shape (D5, D7). Both processes depend on the type of plantation (sun or shade-grown coffee) (N8, N11), the coffee variety (N7), and the topography of the field (N10, N11). Plant renovation dynamics also affect the planting pattern (N8). In the organic plantation individual plant renovations are more common, as well as the presence of other trees apart from coffee. This also defines more irregular patterns than in the conventional plantation without shade trees, where whole locations are removed and renovated every few years to increase short-term productivity. This reflects on the different degrees of randomization of the trajectories of the harvesters during harvest (Fig.1).

The number of harvesters (D3) per *pante* and the *pante* size (D7) could also determine the general pattern as it affects how fast families can harvest one location during the day and travel to another pante (long-steps in Fig. 1). The number of harvesters changes across the harvesting season following the coffee maturation process (N6). Finally, the number of working hours (D6) or the difficulty of harvesting a specific coffee variety (D4) also affect the trajectory length.

Most of these variables can be directly or indirectly affected by management decisions and ultimately depend on the specific syndrome of production of both plantations (green boxes in Fig. 4). For example, owners specify the starting and ending date of the harvesting season. Owners also decide, mainly with profit-maximizing criteria, on the coffee varieties, the shade management, or the dynamics of plant renovation. As we saw, these factors will indirectly affect the final trajectory of the harvesters. Management can also determine the motion of harvesters by changing the working hours or the number of people by *pante*, or the distance between *pantes*. In this sense, the recorded trajectories of both plantations, their lengths, and patterns are the result of two different shade and management practices within a specific syndrome of production.

## 4. DISCUSSION AND CONCLUSIONS

Coffee is produced under a large variety of management practices, land tenure systems, and scales. In Mexico, one finds community-owned rustic coffee plantations as well as large-scale landlord-owned sun, and sometimes shaded, coffee plantations (Moguel and Toledo, 1999). Here we draw upon the conceptual framework of the “syndromes of production” to evaluate some of the social and ecological dynamics within a given agricultural paradigm (Andow and Hidaka, 1989; Ong and Liao, 2020; Vandermeer and Perfecto, 2012). In particular, we maintain that the two studied plantations (organic and conventional) in the Sonocusco region are part of a general capitalist syndrome of production characterized by large-scale plantations where landlords extract surplus value from the work of permanent and temporary workers and sell their product to conventional or organic markets (López Echeverría, 2006; Ong and Liao, 2020). The distinction in their markets reflects differences in their specific management and ecological practices. The organic plantation has an integrated management system that results in higher associated biodiversity, pest control, and lower plant renovation, but also in better worker’s health conditions by removing pesticides (Soto-Pinto et al., 2000; Jiménez-Soto, 2021). Besides, it values quality over quantity in coffee production, which reduces the plant renovation rate and prioritizes planting high-quality arabica plants.

Our study aimed to understand how distinct agricultural managements in large-scale capitalist plantations drive differences in patterns of human movement that can potentially impact CLR spread in the environment, as well as working conditions. We found that two-state space models can describe harvester’s trajectories well. This is in line with our observations in the field where we differentiate two kinds of movements: when trees have berries, harvesters remain in the same rows or nearby (Collect state: short to medium steps); when no berries are in sight or when they have finished a *pante*, harvesters move to another zone of the *pante* or to another *pante* chosen by the foreman (Search state: medium to long steps). These behaviors are similar to general foraging patterns in other agroecosystems (Reynolds et al., 2018) or to the trajectories generated by deterministic models where a walker visits the closest unvisited site among a collection of point-like sites irregularly distributed in a landscape (Santos et al., 2007). The organic shaded plantation had longer steps both in the Collect state (within the *pante*) and specially in the Search state (relocalizations between *pantes*) in the same amount of time and, according to our qualitative analysis this could be related to *a)* a lower fruit load per tree or low percentage of trees with berries when we took the data or *b)* smaller *pantes* or the number of harvesters per *pante*. This resulted in a higher number of visited areas of different sizes (Fig. 3). The heterogeneity in trajectories in this plantation seems to be related to the presence of other trees, plant renovation dynamics, or even to the topographic context, as also discussed by Hajian-Forooshani and Vandermeer, (2021) and Soto-Pinto et al., (2000).

In this sense, the spatial movements of the harvesters are the result of their interactions with biological and climatic factors and also depend on plantation characteristics and management decisions. This goes along the line of the unifying paradigm of movement ecology first introduced in the context of animals (Nathan et al 2008). For example, ripening can be affected by the regularity of the rainfall during flowering and the temperature (Kath et al., 2021), or by the distribution of shade (Henrique Marquezine Leite et al., 2022). Nevertheless, the owners decide when to end the harvesting season. As was reported by other studies, it is common for the owners to encourage harvesters to stay until the end of the season by increasing the value of the harvest load, or to directly force them to remain in the fields by delaying their payment (Jiménez-Soto, 2021). This promotes harvesting when plants have a low percentage of trees with ripe fruits and low fruit load per tree, setting the scenario for longer daily trajectories with a higher proportion of medium to long steps across the region.

There are several potential socio-ecological implications of the different harvesters’ trajectories on coffee rust dispersal. Here we did not report any data related to CLR or other potent ial pests but we can hypothesize that visiting more trees per day, without the possibility of changing clothing could increase the dispersal and impact of CLR (as observed by Becker and Kranz (1977)). Besides, CLR or other pathogens are also dispersed from plant to plant through water splash or leaf-to-leaf contact, creating localized foci of infection (Vandermeer et al., 2018). Changing *pante* several times (Fig. 1, organic plantation) could connect these foci of infection or bear the pathogens to different parts of the landscape and start new infections. The square units depicted in Fig. 3 could be thought of as a proxy of these foci. In this sense, the spatial heterogeneity in coffee maturation creates fragmented host-landscapes that interact with the movement of harvesters and could increase the CLR or other pathogens’ outbreaks (as suggested by White et al., 2018). Nonetheless, these hypotheses need to be tested, theoretically or empirically. These implications would not mean promoting managements that could homogenize the coffee ripening or fruit charge (e.g., homogenous sun coffee varieties, controlled irrigations) to reduce the spatial movement of harvesters during the harvesting. Coffee production and the incidence of specific pathogens as CLR are multifactorial processes (Avelino et al., 2006) and decisions that could reduce human dispersal can promote another dimension of the epidemic. For example, removing the shade trees to avoid irregular ripening (Henrique Marquezine Leite et al., 2022) also increases the CLR wind-mediated dispersal (Gagliardi et al., 2020). They also do not mean placing responsibility on the harvesters as they are temporary or permanent workers paid daily by piecework, working precariously for several hours to maximize the daily harvest in landlord-owned plantations guided by the generation of profit.

Our findings call for organic coffee management that could prevent diseases, increase diversity, and guarantee just and safe conditions for workers. In particular, to reduce the possible impact of human dispersal of pathogens during harvesting, we suggest, among other ideas, the need for shorter trajectories by working fewer hours a day, avoiding harvesting at the beginning or end of the coffee maturation season when few trees have berries and harvesters have to travel longer distances, or even skipping infected plants. These measures can reduce short-term productivity but sustain long-term productivity. In this sense, to be implemented, coffee farmers should overcome the business-as-usual politics of large-scale capitalist coffee agroecosystems.

## 5. ACKNOWLEDGEMENTS

EMVC is a doctoral student from the Programa de Doctorado en Ciencias Biomédicas, Universidad Nacional Autónoma de México, and has received CONACyT scholarship 686776. MB acknowledges financial support from UNAM-DGAPA-PAPIIT (IN207819). EMVC, CGG and MB thanks Gustavo Bautista, Gabriel Domínguez and Elisa Lotero for the discussions and precious help during the fieldwork. EMVC especially thanks Elisa Lotero for her patience and support and the people from La Parcela for their ideas and friendship. Finally the authors dedicate this work to all the harvester families in both plantations and in other regions, that feed the world under social conditions that must be overturned .

