## Supplemental material 1 for "Spatial trajectories of coffee harvesting in large-scale plantations: ecological and management drivers and implications"

### Supplementary Material 1. HMM analysis from: Spatial trajectories of coffee harvesting in large-scale plantations: ecological and management drivers and implications

Mora Van Cauwelaert et al., 2023

#### 0. Introduction to State-space models with HMM

State space models coupled with hidden Markov models (HMMs) are models in which the distribution that generates an observation $Z_t$ depends on the state $S_t$ of an underlying and unobserved Markov process (Fig.1) (Zucchini et al., 2016). In this sense, the observations $Z_t$ can be retrieved from one to multiple distributions related to the number of states. In the context of animal movement, the state $S_t$ is interpreted as a proxy for the behavioral state of the animal (e.g. foraging, exploring; (Michelot et al., 2016)). The observations $Zt$ are bivariate time series ($Zt = (l_t, \phi_t)$) where $l_t$ is the step length (the Euclidean distance) and $\phi_t$ the turning angle, between two succesive locations (Zucchini et al., 2016). Biologically speaking, these models try to include the process where the movement of agents (e.g. short or large steps) depends on its behavior (Patterson et al., 2008). With these models, we can define the different originary distributions and then decode the most likely sequence of states along the trajectory of an agent, its average time within a state, and the number of switches between states.

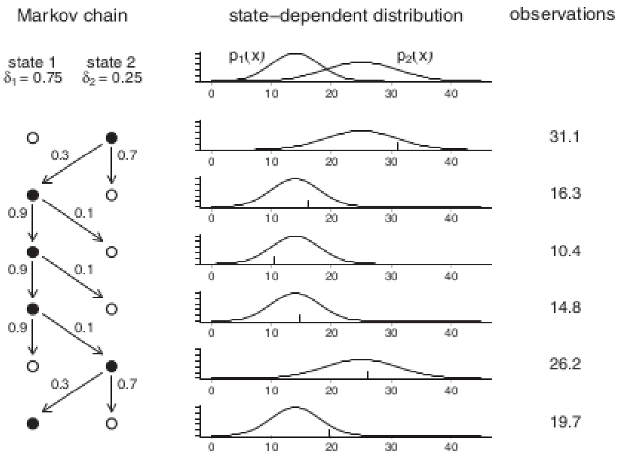


Figure 1: Process generating the observations in a two-state HMM. The chain followed the path 2, 1, 1, 1, 2, 1, as indicated on the left. The corresponding state-dependent distributions are shown in the middle. The observations are generated from the corresponding active distributions. Taken from (Zucchini, 2016)

Here we used the *movehmm* library (Michelot et al., 2019) to i) fit the most likely distributions to the harvester movement data in each plantation, ii) decode and compare the sequence of states in the two plantations (Ecological and Conventional) with the Viterbi algorithm. To see the full results and visualizations please refer to main text.

### 1. Preparation of data and algorithm

#### 1.1. Preparation of data

We loaded *ggplot2*, *dplyr*, *tidyverse* and *movehmm* libraries. We then loaded the data of the trajectories and did some punctual modifications to it. We had in total 12 trajectories, with $(x, y)$ coordinates (see Fig. 2).

We took these time irregular trajectories (where each point represents one different tree, but where the time between two trees is variable) and treated them as regular trajectories. This aimed to analyze the change in the movement of the workers during a day, generated by the underlying pattern of trees or by the differences in the fruit charge and ripening synchronicity. We convert this database into a *movehmm* object where the step distance $l_t$ and the relative angle $\phi_t$ between the $(x, y)$ coordinates is calculated. Now, for the following analysis, we only took the step distance and treated each plantation separately. We decided to exclude the angles because they didn't have a biological meaning for irregular trajectories.

#### 1.2. Fitting algoritm

The algorithm of the *movehmm* library uses:

a) a predefined number of states. Here we will use one and two.

b) a defined family of distributions. Here we will try to fit two different exponential families (*gamma* and *weibull*).

c) prior parameters for each of the distributions, for each state.

d) The observations (step-lengths) (the data of harvesters)

With these inputs, the algorithm extracts the most likely two-state and one-state distributions with their respective parameters (within the predefined family of distributions) from the data. During the fitting process, the algorithm uses the maximum-log likelihood and the *forward algorithm* (a recursive algorithm starting with the prior distributions; (Zucchini et al., 2016)). We ran a loop across 1000 prior parameters within the range shown in Table 1 to fit the data to two different families (*weibull* and *gamma*) and to avoid local maximal likelihoods (Michelot et al., 2019). For the *gamma* distribution, the minimum and maximum values of each prior parameter range were chosen to encompass the full distribution (including values below and over the mean). For the *weibull* distribution, the maximum shape considered the left-skew shape and the scale was chosen to include the 120m long steps (Table 1). Many of these combinations of prior parameters converged to the same final distributions.

Table 1: Minimun and maximum values of the prior parameters for each distribution

| value | gamma.mean | gamma.sd | weibull.shape | weibull.scale |
|---|---|---|---|---|
| min | 0.1 | 0.1 | 0.1 | 0.1 |
| max | 10.0 | 10.0 | 2.7 | 15.0 |

#### 1.3. Outputs

We first ran the model for each plantation (Organic and Conventional) assuming two-state distributions.The output is stored in /output/dataFrameAnalysis/.

We then ran the model for each plantation (Organic and Conventional) assuming one-state distributions. The output is stored in /output/dataFrameAnalysis/

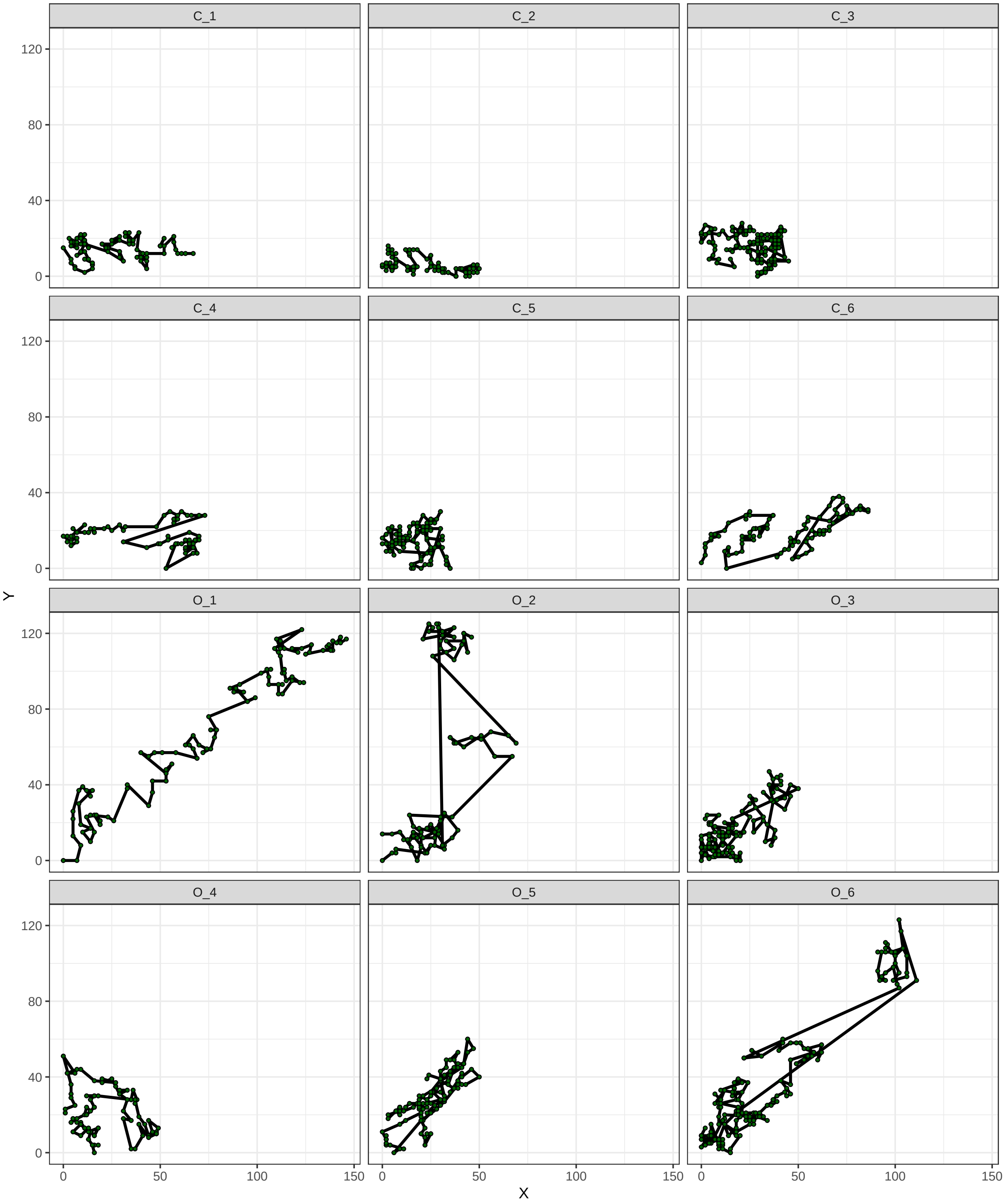


Figure 2: Trajectories of harvesters for both plantations (O: organic, C: conventional). Each green dot represent a tree

#### 2. Results

##### 2.1 Two-state and one-state fitting

Some combinations resulted in distributions with zero variance. We decided to remove those cases as they did not make any biological sense. We also removed states with means higher than the ranges (this does not change the results as they had maximal AIC that would be removed anyways). We then plotted the AIC without these outliers (Fig.3). Almost all the combinations of prior parameters converge to two models with equivalent AIC for both distributions (Fig. 3). In this sense, the parameters of the models are robust to the initial prior parameters chosen. Two-state gamma distributions had the minimal AIC. The minimal AIC for each family and number of states are represented in Table 2 and stored in: output/dataFramesGenerated/

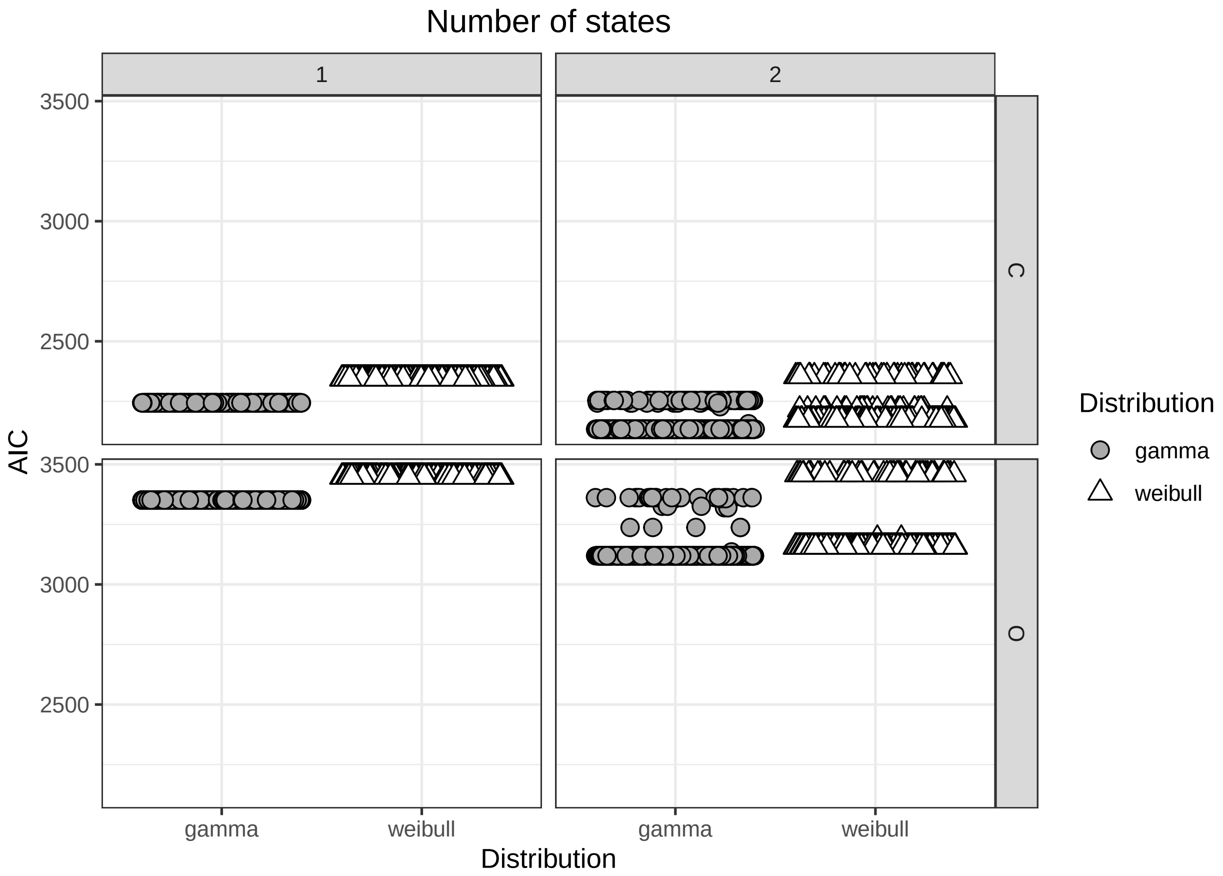


Figure 3: Minimun AIC per combination of prior parameters for two different distributions, for 1 and 2 states, for each of the plantations. O:Organic, C: Conventional

Table 2: Best models for each of the distributions. We added the prior parameters, the min negative likelihood, AIC criteria and the final parameters. pr:prior, st1: state 1, p1: parameter 1 (shape for weibull and mean for gamma), p2: parameter 2 (scale for weibull and sd for gamma).

| model | prior_par0_st1 | prior_par1_st1 | minNegLike | AIC_model | st1_par0 | st1_par1 | farm | prior_par0_st2 | prior_par1_st2 | st2_par0 | st2_par1 | states |
|---|---|---|---|---|---|---|---|---|---|---|---|---|
| weibull | 2.26 | 9.99 | 1169.82 | 2343.64 | 1.47 | 4.66 | C | NA | NA | NA | NA | 1 |
| gamma | 7.79 | 7.35 | 1119.84 | 2243.68 | 4.17 | 2.47 | C | NA | NA | NA | NA | 1 |
| weibull | 1.08 | 2.57 | 1721.75 | 3447.51 | 1.16 | 6.35 | O | NA | NA | NA | NA | 1 |
| gamma | 3.77 | 9.08 | 1673.67 | 3351.34 | 5.94 | 4.29 | O | NA | NA | NA | NA | 1 |
| weibull | 0.14 | 13.68 | 1079.15 | 2172.30 | 1.46 | 10.12 | C | 1.39 | 7.67 | 2.42 | 3.87 | 2 |
| gamma | 8.01 | 6.10 | 1059.92 | 2133.84 | 3.47 | 1.56 | C | 8.49 | 8.06 | 10.03 | 6.41 | 2 |
| weibull | 1.44 | 9.92 | 1571.22 | 3156.44 | 1.96 | 5.47 | O | 0.86 | 13.83 | 0.97 | 21.25 | 2 |
| gamma | 7.89 | 4.90 | 1552.71 | 3119.43 | 5.01 | 2.80 | O | 0.48 | 8.46 | 32.95 | 30.91 | 2 |

##### 2.2 Sequence of states for each plantation (two states gamma distribution)

In the following analysis we only used the two-states gamma distribution as it had the lower AIC. For each plantation we assigned each step to "state 1" or "state 2" according to the Viterbi algorithm. The output tables used for the plotting the results are exported to data/

With the Viterbi algorithm, the package *movehmm* decodes the most likely sequence of states (assuming a markov chain) and the transitions matrix between hidden states. This takes into account the conditional probabilities between the observations and hidden states $P(Z_t|S_t))$ (Zucchini et al., 2016) (Fig. 4 and 5). In this sense, *movehmm* can estimate the state with the highest probability for each step but this might not be the same as the state in the most probable sequence returned by the Viterbi algorithm (Fig. 4 and 5, second and third row vs first row). This is because the Viterbi algorithm performs "global decoding", whereas the state probabilities are "local decoding" (Zucchini et al., 2016). In the main text we plot the result of the Viterbi algoritm for each of the algoritms.

```
## Decoding states sequence... DONE
## Computing states probabilities... DONE
```

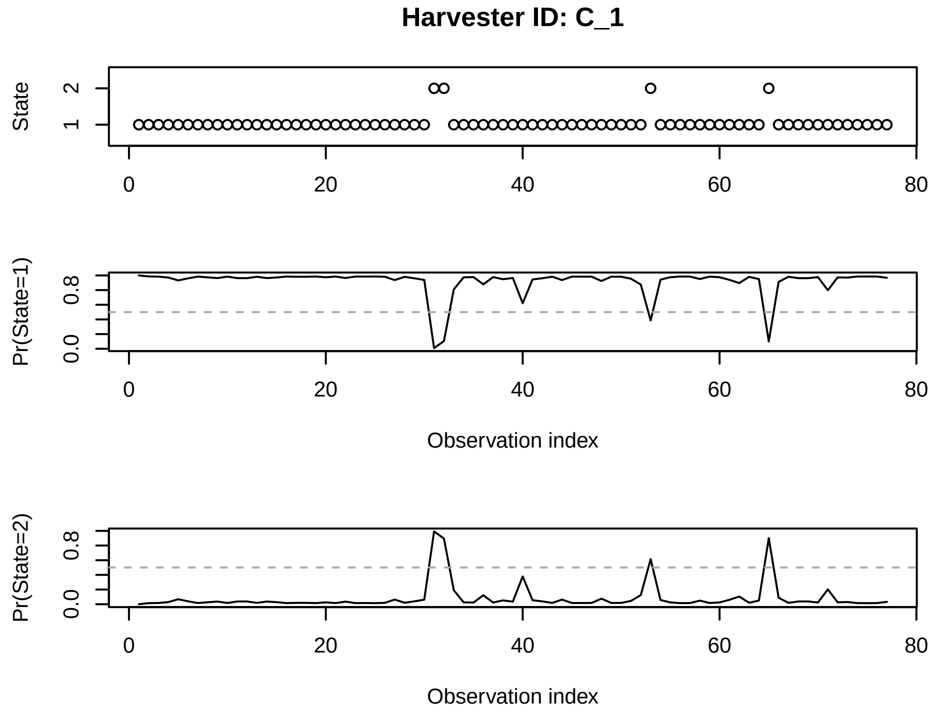


Figure 4: State probabilities for each step of harvester C_1 trajectory and result of the Viterbi algorithm

```
## Decoding states sequence... DONE
## Computing states probabilities... DONE
```

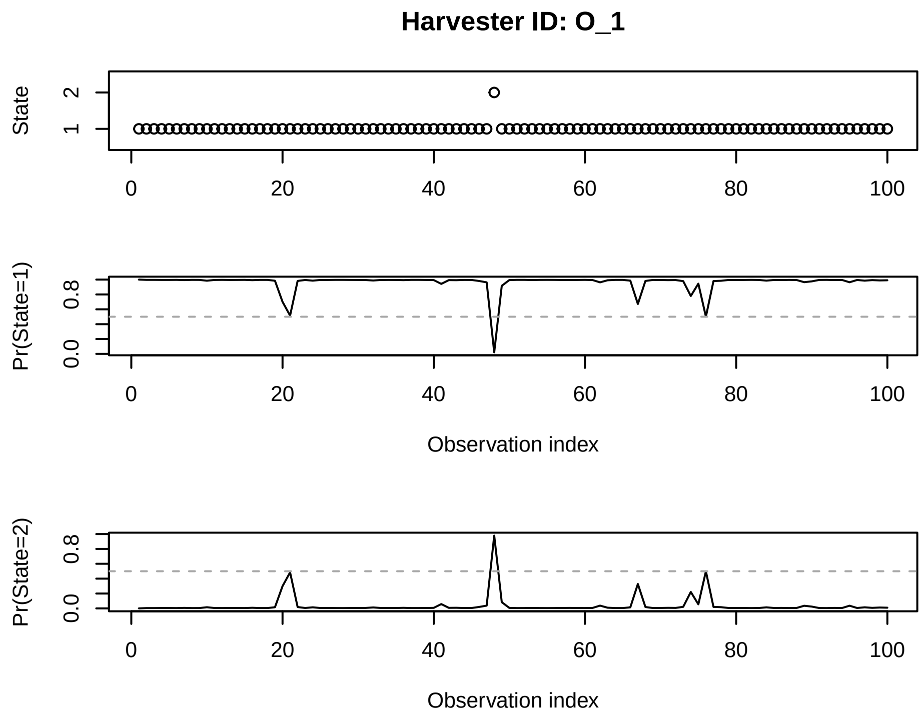


Figure 5: State probabilities for each step of harvester O_1 trajectory and result of the Viterbi algorithm
